# Whole-Brain Vasculature Reconstruction at the Single Capillary Level

**DOI:** 10.1101/191502

**Authors:** Antonino Paolo Di Giovanna, Alessandro Tibo, Ludovico Silvestri, Marie Caroline Müllenbroich, Irene Costantini, Anna Letizia Allegra Mascaro, Leonardo Sacconi, Paolo Frasconi, Francesco Saverio Pavone

## Abstract

The distinct organization of the brain’s vascular network ensures that it is adequately supplied with oxygen and nutrients. However, despite this fundamental role, a detailed reconstruction of the brain-wide vasculature at the capillary level remains elusive, due to insufficient image quality using the best available techniques. Here, we demonstrate a novel approach that improves vascular demarcation by combining CLARITY with a vascular staining approach that can fill the entire blood vessel lumen and imaging with light-sheet fluorescence microscopy. This method significantly improves image contrast, particularly in depth, thereby allowing reliable application of automatic segmentation algorithms, which play an increasingly important role in high-throughput imaging of the terabyte-sized datasets now routinely produced. Furthermore, our novel method is compatible with endogenous fluorescence, thus allowing simultaneous investigations of vasculature and genetically targeted neurons. We believe our new method will be valuable for future brain-wide investigations of the capillary network.

## INTRODUCTION

Neuronal activity relies on an intricate network of blood vessels that delivers oxygen and nutrients for neuronal metabolism. Despite the fundamental importance of this system, we do not have a complete topological understanding of the capillary network through which the exchange of substances and metabolites takes place.

Mapping the fine anatomical details of capillaries over the entire mouse brain has been challenging because it requires micrometre resolution coupled with fast acquisition speeds in order to cover the entire sample volume in a reasonable time period. Although large vessels in the whole mouse brain can be detected non-invasively with a variety of techniques^1-4^, these methods provide a rather coarse resolution. Other minimal invasive methodologies allow only poor visualization of smaller superficial vessels, without reaching capillary-level resolution and without including deep regions^5-8^. The capillary network can be visualized at high resolutions via optical microscopy; however, depth is limited when acquiring images from opaque samples. The development of optical clearing methods^9, 10^ has allowed mouse organs, including the brain, to be rendered completely transparent, thus overcoming this limitation. Among the optical approaches available in combination with tissue clearing, light-sheet fluorescence microscopy (LSFM) offers the highest acquisition speed^11^ and is therefore particularly attractive for reconstructing large volumes of tissues that would otherwise take too long to image using point-scanning techniques like confocal microscopy or two-photon fluorescence microscopy (TPFM). Indeed, clearing methodologies coupled with LSFM have been extensively used for imaging fluorescently-labelled components in whole-mouse organs^12-19^, including the visualization of brain vasculature^14, 18, 19^. Imaging of the vasculature network with LSFM yields datasets of 2–3 TB per brain, and automated analysis is thus paramount for extracting meaningful morphological information from this data. However, because of extremely large amounts of data to be processed and the suboptimal image quality, brain-wide vascular network reconstruction at the capillary level has not been achieved yet. On the contrary, vascular network reconstruction has been accomplished only within a portion of mouse brain cortex using TPFM^20^. The outstanding result of this latter work has been made possible by the use of a novel vascular staining method by which the whole blood vessel lumen is filled with a fluorescent gel^21^. The applicability of such a staining method to whole brain acquisitions with LSFM is not straightforward because it needs to be compatible with a whole brain clearing procedure. Very recently, a methodology combining this labelling approach with 3DISCO^18^ whole brain clearing was proposed. However, the corrosive effect of the mounting medium necessitated the use of a specific objective lens with low magnification and coarse resolution.

Here, we present a methodology that combines blood vessel lumen staining^21^ with CLARITY^12^ and high-resolution imaging with a custom-made LSFM apparatus^22^. We chose tissue transformation with CLARITY as it is effective in achieving full transparency and preserves endogenous fluorescence. CLARITY generates tissue transparency through the selective removal of lipids and the retention of proteins via binding of functional groups to acrylamide and PFA monomers. The blood vessel lumen staining we use here is composed of fluorophore albumin-FITC conjugate (BSA-FITC) dissolved in porcine skin gelatine. Because the fluorescent marker and the gel are comprised of proteins, they are both retained in the hydrogel matrix during the aggressive wash-out of the lipids. Additionally, the high molecular weight of albumin prevents the marker from crossing blood vessel walls, which ensures the confinement of the fluorescent signal within the blood vessels.

As a starting point, we demonstrate that staining of the blood-vessel lumen leads to an increased SNR and provides clear demarcation of blood vessels. As proof of the high image quality obtained, a simple automatic segmentation algorithm worked well when applied to a transverse section of the whole-brain vascular dataset acquired with LSFM. Furthermore, we show that our clearing and staining protocol does not introduce morphological changes that could result in inaccurate mapping. Finally, the good preservation of endogenous fluorescence allow us to perform sequential investigations of both vascular and neuronal morphology in the same brain using *Thy1*-GFP-M transgenic mice.

## RESULTS

### BLOOD VESSEL LUMEN STAINING OUTPERFORMS VESSEL WALL STAINING

As a first step, we performed a quantitative comparison of the signal-to-noise ratio (SNR) achieved at different depths using the vessel lumen staining developed by Tsai et al., 2009^21^ (hereafter gel-BSA-FITC) and an extensively used endothelial marker (lectin-FITC), acquiring images with two-photon fluorescence microscopy (TPFM) from 2 mm thick fixed-brain slices. Because in fixed tissues TPFM can image to a maximum depth of approximately 200 μm, we used a mild clearing agent (TDE)^23^ to extend our analysis into deeper regions without any tissue transformation. The labellings resulting from the two methods are visually compared in Figure 1 (a-d). While with lectin-FITC staining only the vessel walls are visible (Fig1a, c), gel-BSA-FITC staining results in fluorescent labelling of the whole space inside blood vessels (Fig.1b, d). As predicted, a greater (SNR) was detected at all depths when fluorescent markers filled the entire lumen than when only the vessel walls were labelled (Fig. 1e). Averaging the SNR across depths, a ten-fold increase in SNR was observed using gel-BSA-FITC (Fig. 1f).

**Figure 1.**
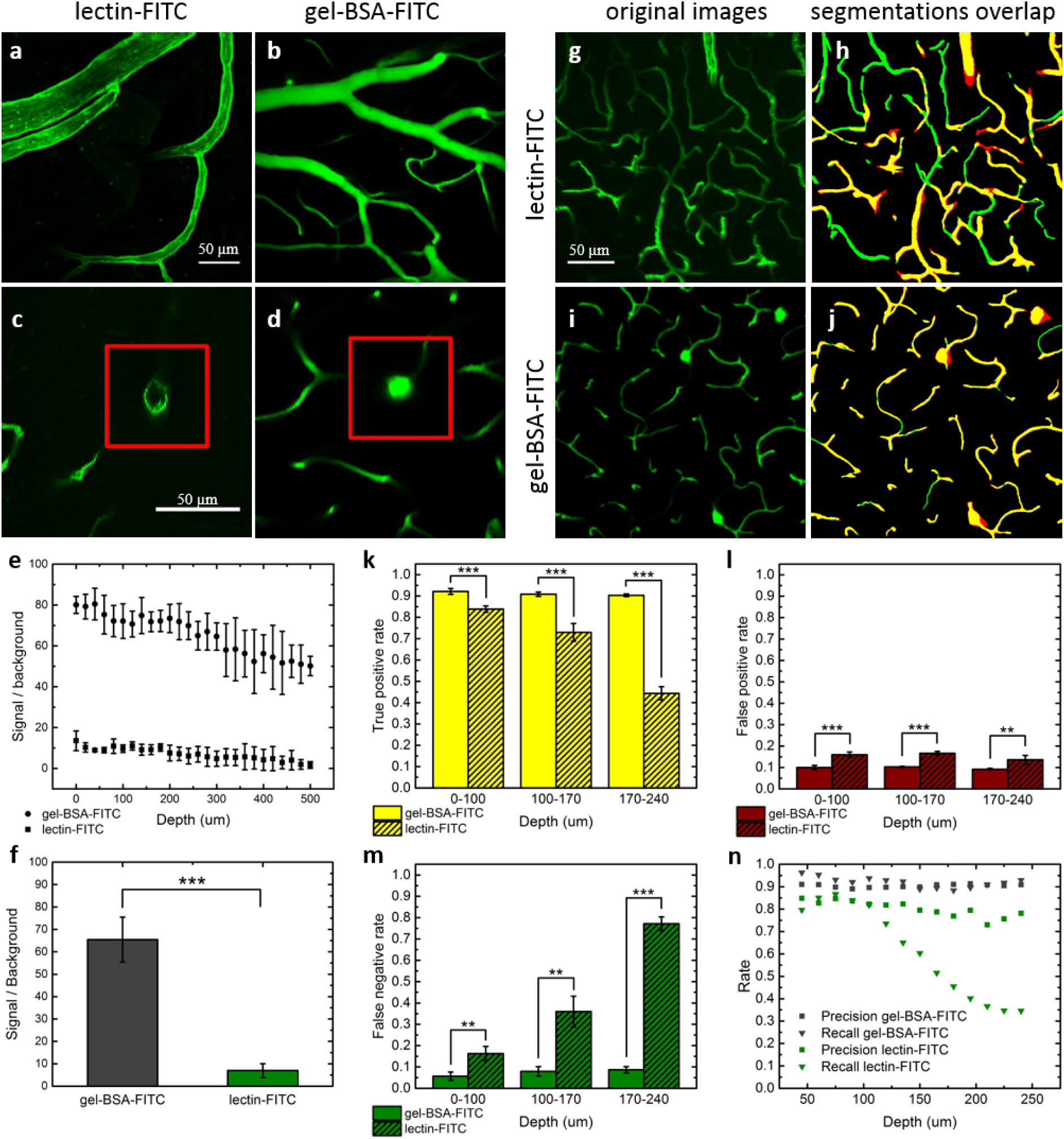
Comparison between lectin-FITC and Gel-BSA-FITC mouse brain vascular staining. (**a, b**) Two-photon acquisition of vessels arranged horizontally with respect to the acquisition plane. (**c**, **d)** Blood vessels arranged perpendicularly to the plane of imaging, enclosed by red insets. Note the absence of fluorescence in the blood vessel lumens in **c**. (**e**) Analysis of the signal to noise ratio by depth (mean ± s.d., n = 6,). (**f**) Average signal to noise ratios across all depths (mean ± s.d.) (***, p < 0.001, One Way ANOVA). (**g**) 40 μm MIP of images from lectin-FITC stained sample acquired from 115 to 155 um of depth with TPFM. (**h**) Overlap between manual (green) and automatic (red) segmentation. (**i**) 40 μm MIP of images from gel-BSA-FITC stained sample acquired from 115 to 155 um of depth with TPFM. (**j**) Overlap between manual (green) and automatic (red) segmentation. (**k**) True positive, (**l**) false positive, and (**m**) false negative rate at different depths for gel-BSA-FITC and lectin-FITC labelling (mean ± s.d., n = 4) (**, p < 0.01, ***, p < 0.001, One Way ANOVA). (**n**) Precision and recall of the segmentations according to depth.

We subsequently verified how the different image quality affected automatic image segmentation. Image stacks acquired from lectin-FITC- and gel-BSA-FITC-stained samples were segmented using an automatic segmentation algorithm based on Markov random fields.^24^ Automatic segmentation accuracy was quantified by superimposing a manual segmentation of the same maximum intensity projections (MIPs) (Fig. 1g-j) and reporting the percentages of shared pixels (true positives), pixels detected only by automatic segmentation (false positives), and those detected only by manual segmentation (false negatives). When using gel-BSA-FITC staining, the true positive rate remained stable at more than 90% over the range of depths inspected (0-240 μm) (Fig. 1k). However, when using lectin-FITC staining, we observed a rapid drop from 84% to 44% at a depth of 240 μm. The false positive rate (i.e., overestimation; algorithm segmentation > manual segmentation) was slightly but significantly higher using lectin-FITC (~10% with gel-BSA-FITC vs ~16% with lectin-FITC for all depths) (Fig. 1l). Owing to the weaker contrast, the false negative rate (underestimation) for lectin-FITC staining increased drastically with imaging depth (100-μm depth: 16%; >200-μm depth: 77%) (Fig. 1m). In contrast, gel-BSA-FITC staining resulted in only a slight increase in underestimation with imaging depth (from 5% to 8%). The counts of true positives (TP), false positives (FP), and false negatives (FN) were used to estimate the precision *P* = TP/(TP + FP) and the recall *R* = TP/(TP + FN) at varying depths (Fig. 1n). Although precision remained stable for both methods over a range of depths, it was higher for the gel-BSA-FITC labelling. While a large drop in recall was observed for lectin-FITC labelling, recall was high and stable with gel-BSA-FITC labelling, even at high imaging depth.

### HIGHLY ACCURATE WHOLE MOUSE-BRAIN VASCULATURE TOMOGRAPHY WITH LSFM

In order to use the gel-BSA-FITC staining in association with CLARITY clearing, we split the staining and clearing into two subsequent passages. This was because the different temperatures needed to polymerize the gelatine and the hydrogel solution used in the CLARITY protocol, make the intracardiac injections of both incompatible. Therefore, we first perfused the mouse with paraformaldehyde (PFA) and gel-BSA-FITC for vascular staining (Fig. 2). After that, we incubated fixed samples in a hydrogel solution for a few days to allow passive diffusion of acrylamide monomers and PFA through the tissue. After the subsequent step of lipids removal, TDE clearing^23^ was applied to the CLARITY-processed brain before imaging. The protocol described was well suited for LSFM imaging and allowed obtaining high-quality whole mouse-brain vasculature datasets, whose volumetric reconstruction (from downsampled data) is shown in figure 3 and supplementary video 1. Due to the extremely large image dataset, a 3D detailed visualization of the capillary network is nowadays not possible for the whole brain, even using powerful computer platforms. 3D visualization on higher resolution (downsampled by factor 2 from original data) is shown for a region of interest (ROI) in supplementary video 2.

**Figure 2.**
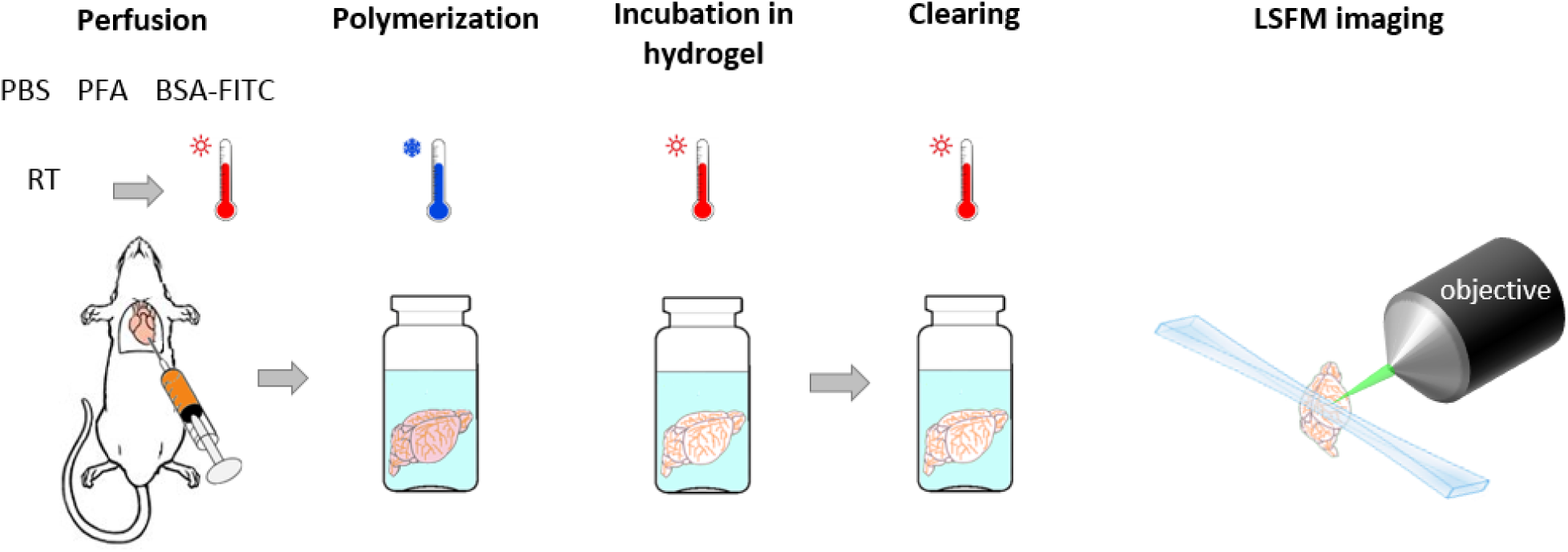
Sample preparation and imaging. The mouse is perfused intracardially with PBS and PFA at room temperature followed by injection of the fluorescent gel kept at 40 °C until injection. Gel polymerization is promoted by placing the carcass in ice cold water soon after gel perfusion. After brain extraction, hydrogel monomers are allowed to diffuse passively through the tissue by incubating the sample in a hydrogel solution for five days. The incubation is followed by tissue clearing with the CLARITY-TDE protocol and imaging with LSFM.

**Figure 3.**
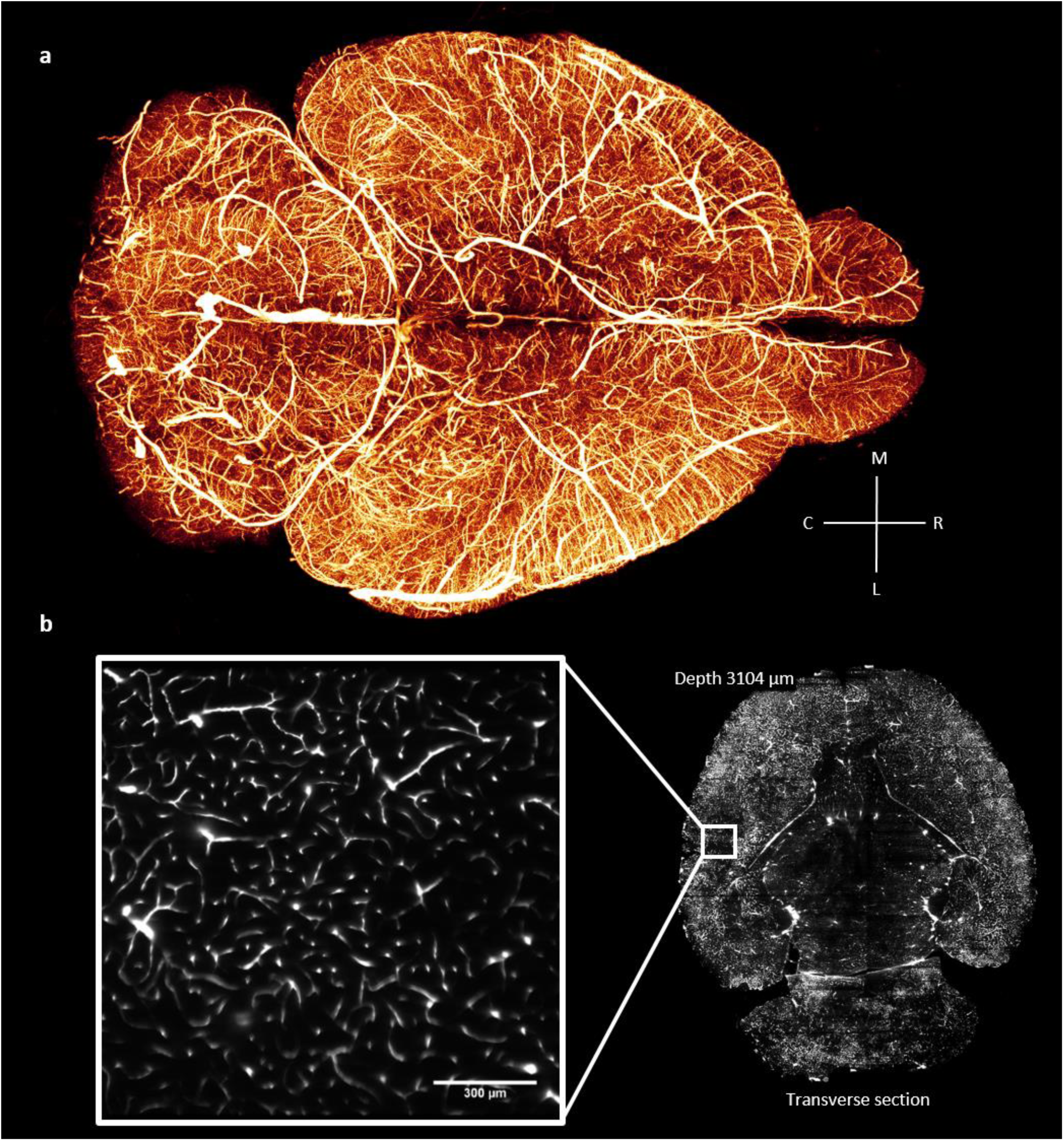
Whole mouse brain-vasculature tomography. (**a**) 3D rendering (Amira 5.3 software, Visage Imaging) of the whole mouse-brain vasculature acquired using LSFM and gel-BSA-FITC vascular staining. The image was produced from downsampled stacks stitched using TeraStitcher software^38^ (**b**) single frame from a stack at original resolution (pixel size 0.65 μm) showing details at the capillary level in a very internal optical section of the brain.

To assess any topological artefacts that could result from morphological changes brought on by the staining or clearing procedure, we performed a correlation analysis at the micrometre level using TPFM and LSFM on the same brain region^25^. *In vivo* TPFM was conducted through a cranial window implanted on a mouse (Fig. 4a) after injection of the fluorescent marker Texas red-Dextran into the blood stream. After vascular staining with gel-BSA-FITC and brain extraction, the same area was later identified and imaged *ex vivo* with TPFM (Fig. 4b). Finally, after CLARITY-TDE clearing, the entire brain was imaged with LSFM and the same region acquired *in vivo* and *ex vivo* using TPFM was retrieved from the whole-brain dataset and the corresponding LSFM image was used for morphological comparison (Fig. 4c). The general morphology appeared to be well preserved according to visual inspection, confirming that the fluorescent gel was appropriately integrated and retained in the hydrogel mesh. We therefore performed a detailed quantitative analysis of vessel diameter and vessel length, for each of the three imaging methods (Fig. 4d-g). When comparing the *in vivo* and *ex vivo* TPFM blood vessel diameter, the slope of the fitted curve showed a general increase in vessel diameter for *ex vivo* TPFM. However, the majority of smaller capillary segments were above the fitted curve, which suggests a different trend for small vessels, specifically capillary shrinkage. The differences between *in vivo* and *ex vivo* vessel diameter before clearing can be ascribed to altered flow rates during perfusion along with the effects of the fixative solution (PFA). The same measurements were performed on LSFM images obtained after CLARITY-TDE clearing of the whole brain. The clearing procedure itself did not introduce further alterations in vessel diameter, as evidenced by the close slope values found for *ex vivo* TPFM and *ex vivo* CLARITY-TDE measurements when correlated with *in vivo* measurements (Fig. 4e).

**Figure 4.**
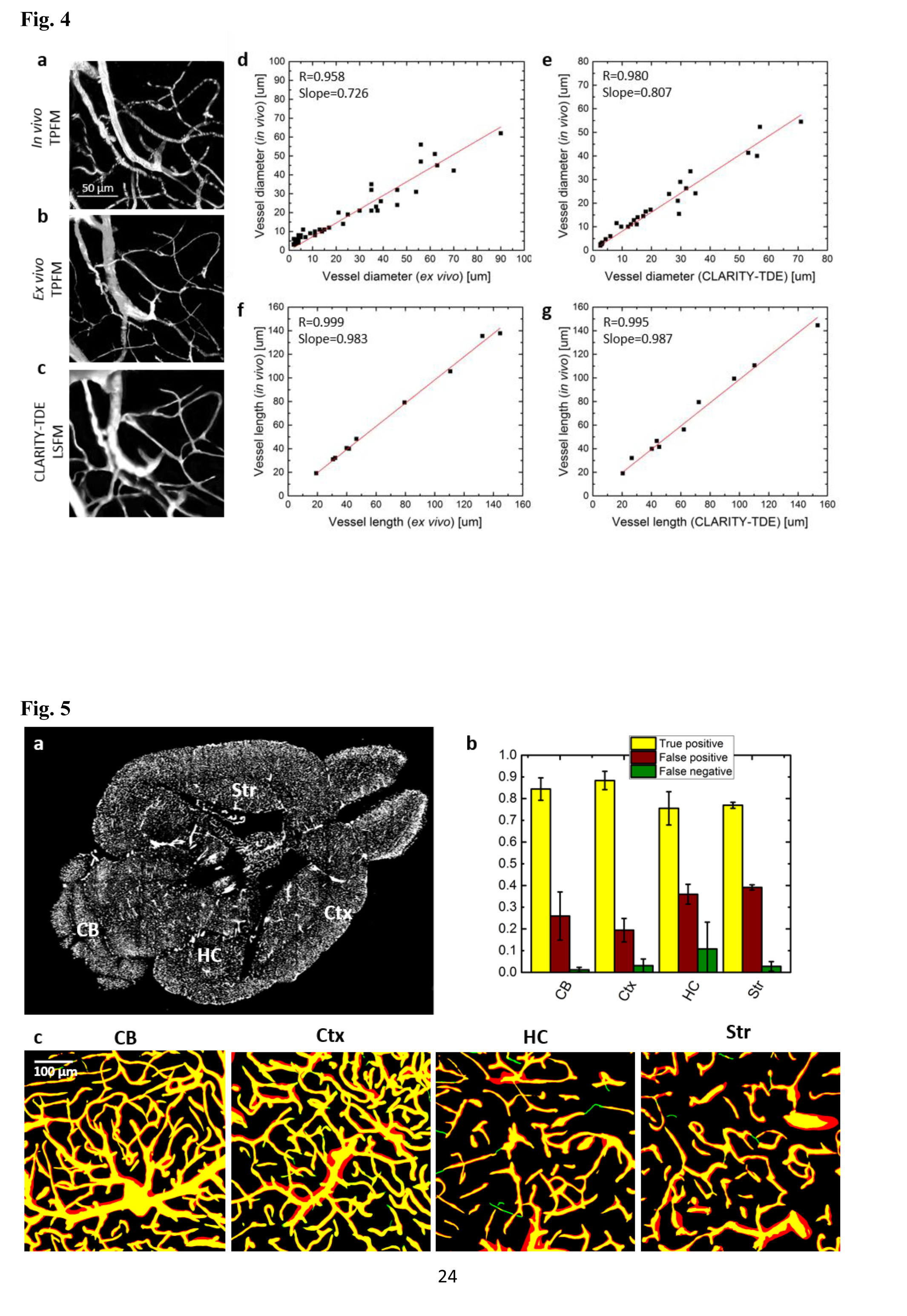
Assessment of morphological changes during brain preparation. **(a-c)** A single mouse brain region of the motor cortex imaged using two-photon fluorescence microscopy (TPFM) *in vivo* through a cranial window (Texas-red Dextran labelling) (**a**), *ex vivo* TPFM (Gel-BSA-FITC labelling (**b**), and light-sheet fluorescence microscopy (LSFM) after tissue clearing (**c**). (**d, e**) Comparison of vessel dimensions between *in vivo* and *ex vivo* TPFM (**d**) and between *in vivo* TPFM and LSFM on clarified sample (**e**). (**f**, **g**) Comparison of vessel segment lengths between *in vivo* and *ex vivo* TPFM (**f**) and between *in vivo* TPFM and LSFM on clarified sample (**g**).

Vessel-segment length measured in three dimensions was compared across the three imaging modalities. We observed that the staining and clearing procedures did not induce any significant additional changes to the vascular structure, as evidenced by the slopes that neared unity (Fig. 4f, g). Overall, very high correlation values (R = 0.999 for *ex vivo* TPFM, R = 0.995 for CLARITY-TDE) indicated good preservation at the single vessel level, proving that vascular tracing is reliable with the proposed approach.

### AUTOMATIC VASCULATURE SEGMENTATION USING THE WHOLE-BRAIN DATASET

Whole-brain data is often several terabytes in size, and analysing it requires automatic image-segmentation algorithms to keep pace with the huge data load. Although complex image segmentation algorithms based on machine learning strategies certainly represent the best choice for yielding the most accurate data, they are time consuming and do not scale easily with high-throughput whole-brain imaging.

Here, thanks to the excellent blood-vessel discrimination obtained with our protocol, we successfully applied a simple and fast unsupervised segmentation method in different regions of an internal section of the mouse brain (Fig. 5a) and evaluated the true positive, false positive, false negative rates with respect to a manual segmentation of the same ROIs (Fig. 5b, c). The true positive values ranged from around 90% in the cerebral cortex to 75% in the hippocampus. The apparently high false positive rate (ranging from 25% to 39 %) was due to different estimations of vessel diameter between the automated and manual segmentations and did not indicate additional vessel branches. For most of the ROIs investigated, the false negative rate was relatively low – usually around 3%, although for two of the selected ROIs in the hippocampus it was 10% and 25%. Notably, the automatic segmentation method was based on a Markov random field^24^, which is an unsupervised segmentation algorithm. Moreover, the images processed belonged to an internal optical section of the brain dataset, which highlights the good image quality achieved throughout the whole sample.

**Figure 5.**
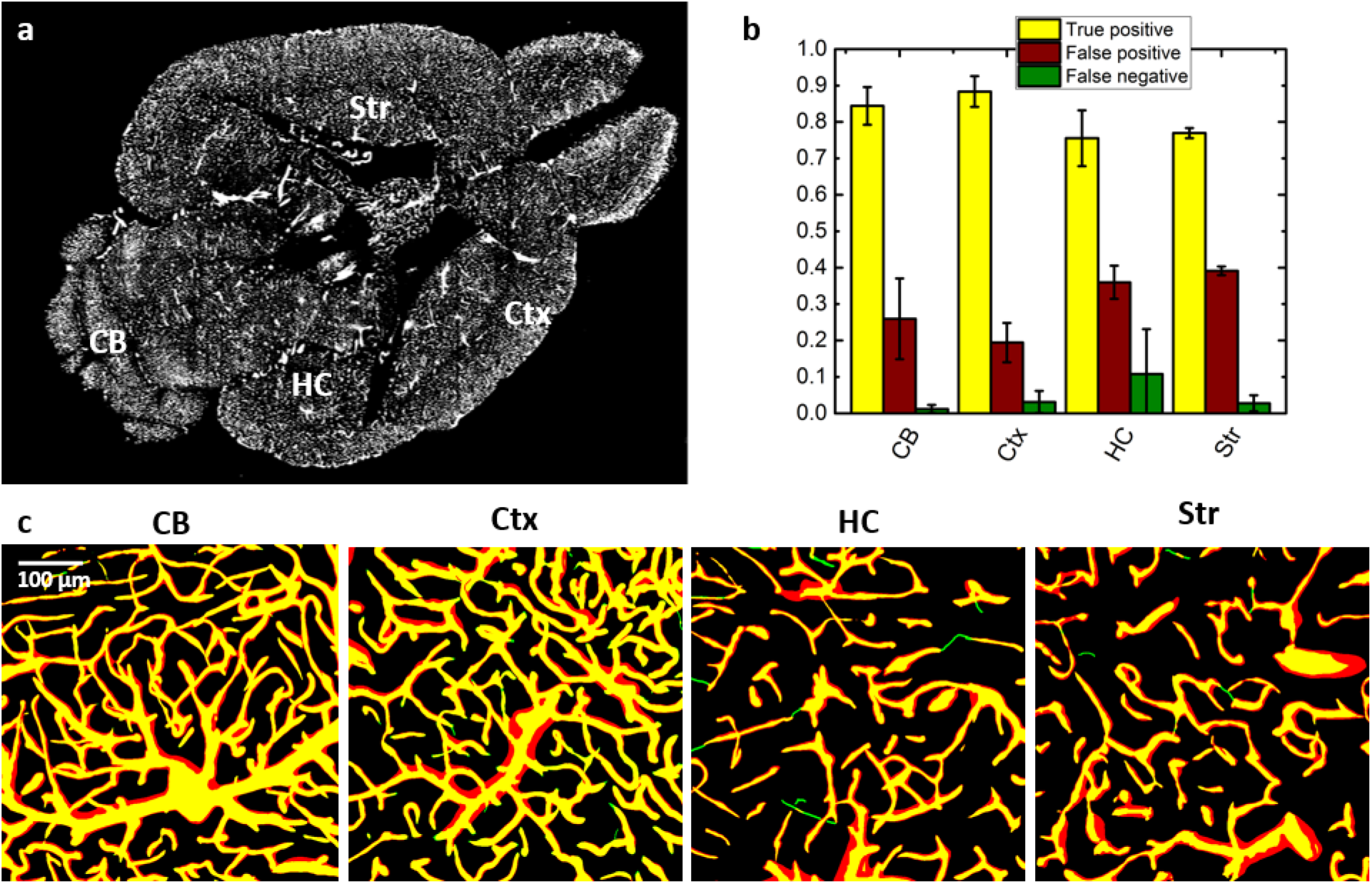
Vasculature segmentation from LSFM images. **(a)** Transverse section of the whole mouse brain acquired with LSFM. The labelled regions are cortex (Ctx), cerebellum (CB), hippocampus (HC), and striatum (Str). (**b**) Quantitative analysis of true positive, false positive, and false negative pixel rates of the automatic segmentation estimated from the overlap between automatically and manually segmentated ROIs from high resolution stacks (mean ± s.d., n = 4) in different brain regions. (**c**) Overlap between manual (green) and automatic (red) segmentation of 40 μm MIPs from different ROIs. Overlapping vessels are showed in yellow.

### VASCULAR AND NEURONAL IMAGING ON *THY1*-GFP-M MICE

Since CLARITY preserves endogenous fluorescence well, we explored the possibility to combine brain-wide blood vessel lumen imaging with neuronal imaging using the *Thy1*-GFP-M mouse line, in which GFP is expressed in a subset of projection neurons^26^. Because of the spectral overlap between GFP and FITC, a different fluorophore—BSA-TRITC conjugate—was chosen to replace BSA-FITC. We determined whether our vessel-staining and clearing procedure was compatible with LSFM imaging of GFP-expressing neurons. We found that the protocol did not affect LSFM neuronal imaging and demonstrated the possibility to perform whole-brain vascular and neuronal imaging of the same sample (Fig. 6 and supplementary video 3 and 4). We highlight that this result represents the first report in which both blood-vessel lumen and GFP-labelled neurons were imaged from the same brain. This prospect expands the applicability of our staining method to include correlation analyses between vascular and neuronal network.

**Figure 6.**
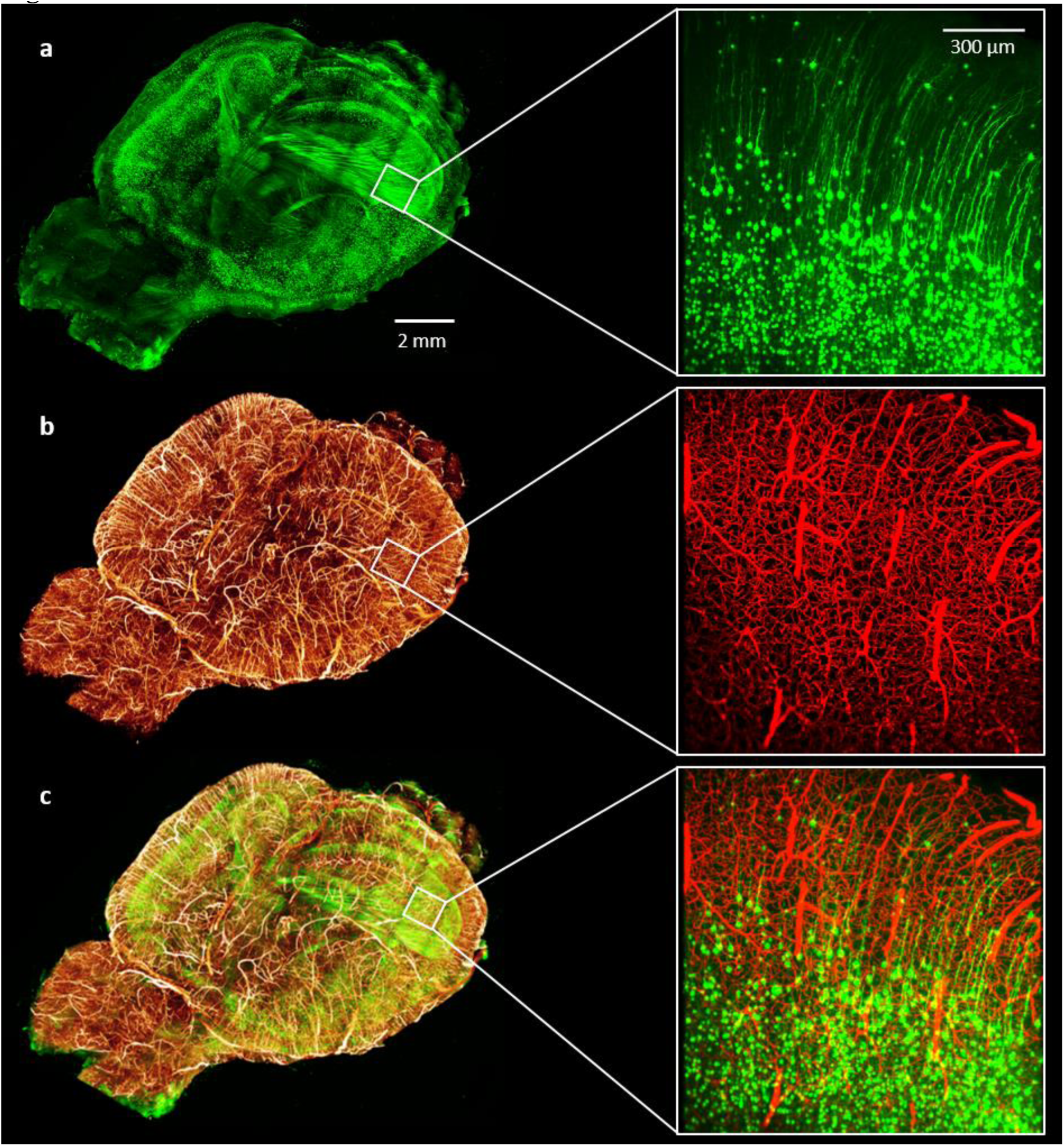
Whole mouse brain neuronal and vascular imaging. (left) **(a)** Whole-brain 3D rendering of GFP-expressing neurons acquired with LSFM. (**b**) 3D rendering of the same brain showing blood vessels labeled with gel-BSA-FITC acquired with LSFM. DownsampleStacks from LSFM acquisitions were stitched using TeraStitcher software^38^. (**c**) Composite image. (right) 100 um MIPs from original acquisitions (*z*-step 2 um, *xy* pixel size 0.65 μm). (**a**) GFP-expressing neurons; (**b**) gel-BSA-FITC labelled vasculature; (**c**) composite.

## DISCUSSION

With the presented article, we describe a new approach for investigations of the vascular network in intact whole mouse-brains. We propose this new method with the perspective of yielding better whole-brain datasets in term of image quality and resolution, as a step forward towards the achievement of whole-brain vasculature network reconstruction. We achieved high quality imaging coupling vessel lumen staining with CLARITY-TDE clearing and applying LSFM. The higher fluorescent signal detected with respect to vessel wall staining, facilitated the identification of blood vessels from the surrounding space by automatic tools, especially in the most internal regions. Manual analysis of morphological alterations with respect to *in vivo* imaging showed some alterations in blood vessel diameter. Such morphological changes were present *ex vivo* before clearing, and did not increase further following the clearing step. Hence, the coupling of the vessel lumen staining with CLARITY-TDE does not *per se* cause alterations, those being instead rather a consequence of the fixation and/or labelling steps. Although some small variations in blood vessel diameter were introduced with respect to the *in vivo* measurements, it is important to consider the effects of neurovascular coupling: regulated cerebral blood flow in living animals produces differential perfusion of brain areas based on cerebral activity and determines changes of vessel intraluminal diameter *in vivo* ^27^. That said, blood vessel diameter reflects specific conditions of brain activity and must be investigated *in vivo* for accurate analysis. Conversely, the length of vessel segments was not altered with respect to *in vivo* measurements. Therefore, the overall topology was well maintained, making the method reliable for software-based reconstruction of the vascular network by means of tracing tools.

Analysis of whole-brain data, which is often several terabytes in size, requires automatic processing to keep pace with the huge data load. However, poor image quality crucially affects the success of automated identification of blood vessels via image segmentation algorithms. This issue can be partly faced computationally, as demonstrated by the development of sophisticated algorithms that are able to identify and fix segmentation artefacts^28^. Additionally, supervised machine learning strategies have been used for automatic identification of brain structures^29^ and can help overcome staining inhomogeneities. Such methods require not only significant computational resources at the time of prediction (thus making segmentation of the whole-brain dataset a demanding process), but also carefully manually-segmented regions as the ground truth used for supervision. By applying an optical solution to the problem, our novel method yield high quality images that can then be processed with computationally simple algorithms, thus reducing the time required for data processing as well as the amount of required training data (and therefore the amount of time required for human experts to perform manual segmentation). Although automatic segmentation from LSFM acquisition was quite good, some differences with respect to the manual segmentation were found and we infer that the use of more sophisticated algorithms will yield more accurate results. We believe that in conjunction with future developments in information processing, our proposed method will enable reliable brain-wide datasets to be obtained, and ultimately improve our understanding of the brain vascular system.

The applications span from studies of morphological variability in physiological conditions to the identification of cerebral vascular phenotypes associate with specific pathologies in which vascular alteration occurs, such as cancer^30^, ischemia^31^ and Alzheimer’s disease^32^. Moreover, the full compatibility with endogenous fluorescence opens up the possibility to investigate the relationship between neuronal and vascular networks on a whole-brain scale using transgenic animals. Evidences indicate that neurons and blood vessels interplay between each other during the wiring processes^33^. Investigations involving both blood vessels and neurons might provide new clues about the mechanisms occurring during the regeneration process after brain damages.

We further anticipate that vascular analysis with this approach could be carried out to create a map that can be superimposed to BOLD-fMRI acquisitions. *In vivo* measurements of microvascular geometry have been previously performed with TPFM with the aim of localizing the BOLD signal detected^34^. The possibility to analyse vascular morphology in broader regions with LSFM will be beneficial for a correct interpretation of data gained from BOLD-fMRI.

The reported examples regarding possible applications suggest that the improved vascular imaging method presented here will be useful across different research groups. Offering new prospects, and addressing a broad audience, the methodology presented will soon have a strong impact on brain research. In addition, we point out that the vascular staining is not restricted to the brain, since all the body is perfused with the fluorescent gel. With the appropriate changes on the clearing procedure, such combination between vessel lumen staining and CLARITY tissue clearing can be optimized for analysis of other organs, expanding further the research fields which could benefit of such vascular investigation strategy.

## METHODS

### Animal models

Adult mice from the C57 line were used for blood vessels tomographies. For neuronal and blood vessels imaging of the same brain, a transgenic line (*Thy1*-GFP-M line) of mice expressing the Green Fluorescent Protein (GFP) in sparse pyramidal neurons was used^35^. All experimental protocols involving animals were designed in accordance with the laws of the Italian Ministry of Health

### Staining the blood-vessel lumen

We used the protocol described in Tsai et al.^21^ for staining the blood-vessel lumen. After deep anaesthesia with isoflurane inhalation, mice were transcardially perfused first with 20–30 ml of 0.01M phosphate buffered saline (PBS) solution (pH 7.6) and then with 60 ml of 4% (w/v) paraformaldehyde (PFA) in PBS. This was followed by perfusion with 10 ml of a fluorescent gel perfusate with the body of the mouse tilted 30° head down to ensure that the large surface vessels remained filled with the gel perfusate. The body of the mouse was submerged in ice water and the heart was clamped to rapidly cool and solidify the gel as the final portion of the gel perfusate was pushed through. After 30 min of cooling, the brain was carefully extracted to avoid damage to pial vessels and then incubated overnight in 4% PFA in PBS at 4°C. The following day, the brain was rinsed three times in PBS.

### Gel compositions

Gel solutions were made of porcine skin gelatine type A (no. G1890; Sigma-Aldrich, Saint Louis, Missouri, USA) in which fluorescein (FITC)-conjugated albumin (no. A9771; Sigma-Aldrich, Saint Louis, Missouri, USA) or Tetramethylrhodamine (TRITC)-conjugate albumin (no. A23016; Waltham, Massachusetts, USA) were dissolved. A 2% (w/v) solution of gelatine was prepared in boiling PBS and allowed to cool to <50°C. It was then combined with 1% (w/v) albumin-FITC (BSA-FITC) or 0.05% albumin-TRITC (BSA-TRITC). The solution was kept at 40°C with stirring before perfusion.

### Vessel staining with Lectin-FITC

For staining the blood-vessel walls, mice were anesthetized with isoflurane and manually perfused with 15 ml of ice-cold PBS 0.01M. Mice were then tilted 30° head down and perfused with 10 ml of ice-cold PBS containing 0.1 mg/ml lectin-FITC from Lycopersicon esculentum (no. L0401; Sigma-Aldrich, Saint Louis, Missouri, USA). After 7 min of incubation, during which lectins were allowed to firmly bind the vessel wall, 40 ml of 4% PFA in PBS was injected. The brain was extracted and incubated overnight in 4% PFA in PBS at 4°C. After incubation, the brain was rinsed three times in PBS.

### Tissue transformation with CLARITY

Fixed mouse brains were incubated in Hydrogel solution (4% [wt/vol] PFA, 4% [wt/vol] acrylamide, 0.05% [wt/vol] bisacrilamide, 0.25% [wt/vol] VA044) in 0.01M PBS at 4°C for 5 days. Samples were degassed and incubated at 37°C for 3 hours to allow hydrogel polymerization. Subsequently, brains were extracted from the polymerized gel and incubated in clearing solution (sodium borate buffer 200 mM, 4% [wt/vol] sodium dodecyl sulfate, pH 8.5) at 37°C for one month while gently shaking. The samples were then washed with PBST (0.1% TritonX in 1X PBS) twice for 24 hours each at room temperature. The brains were then cleared following TDE clearing before imaging.

### TDE clearing

For two-photon imaging of 2-mm thick fixed-brain slices labelled with gel-BSA-FITC or lectin-FITC, the samples were cleared with serial incubations in 20 ml of 20% and 47% (vol/vol) 2,2'-thiodiethanol in 0.01M PBS (TDE/PBS), each for 1 hour at 37°C in gentle oscillation.

For LSFM imaging, CLARITY-processed mouse brains were cleared with 50 ml of 30% TDE/PBS for one day followed by two days of incubation in 63% TDE/PBS, both at 37°C in gentle oscillation. After TDE clearing, CLARITY-processed brains were used for LSFM imaging.

### Two-Photon Fluorescence Microscopy Imaging

A custom-made two-photon fluorescence microscope (TPFM) was assembled from a mode-locked Ti:Sapphire laser (Chameleon, 120 fs pulse width, 80 MHz repetition rate, Coherent, CA) coupled with a custom-made scanning system based on a pair of galvanometric mirrors (VM500+, Cambridge Technologies, MA). The laser light was focused onto the specimen using a water-immersion 20× objective lens (XLUM 20, NA 0.95, WD 2 mm, Olympus, Japan) for fixed specimens and *in vivo* measurements. For Gel-BSA-FITC and lectin-FITC stained samples cleared with 47% TDE/PBS (RI 1.42), we used a tuneable 20× objective lens (Sca/e LD SC Plan-Apochromat, NA 1, WD 5.6 mm, Zeiss, Germany). Imaging in depth was performed with 2-μm z-step. The system was equipped with a motorized *xy* stage (MPC-200, Sutter Instrumente, CA) for lateral displacement of the sample and with a closed-loop piezoelectric stage (ND72Z2LAQ PIFOC objective scanning system, 2-mm travel range, Physik Instrumente, Germany) for the displacement of the objective along the *z* axis. The fluorescent light was separated from the laser optical path by a dichroic beam splitter (DM1) positioned as close as possible to the objective lens (non-de-scanning mode). A two-photon fluorescence cut-off filter (720 SP) eliminated rejected laser light. A second dichroic mirror (DM2) was used to split the two spectral components of the fluorescence signal. The fluorescence signals were filtered with 630/69 and 510/42 filters (FF1 and FF2) and collected by two orthogonal photomultiplier modules (H7422P, Hamamatsu Photonics, Japan). The instrument was controlled by custom software, written in LabView (National Instruments, TX).

### Signal-to-noise ratio measurements

Lectin-FITC or gel-BSA-FITC stained samples were cut into 2-mm thick coronal slices using a vibratome and cleared with 47% TDE/PBS. The signal-to-noise ratio (SNR) was calculated from the images every 20 μm over a depth of 500 μm. The mean grey values of selected ROIs in internal area of vessel or the surrounding area were used as values of signal and background respectively. Six stacks were used for each sample.

### Light-sheet microscopy imaging

Whole brains were imaged using a custom-made light-sheet microscope described in Muellenbroich et al.^22^ The light sheet was generated using a laser beam scanned by a galvanometric mirror (6220H, Cambridge Technology, MA); confocality was achieved by synchronizing the galvo scanner with the line read-out of the sCMOS camera (Orca Flash4.0, Hamamatsu Photonics, Japan). The laser light was provided by a diode laser (Excelsior 488, Spectra Physics) and an acoustooptic tuneable filter (AOTFnC-400.650-TN, AA Opto-Electronic, France) was used to regulate laser power. The excitation wavelengths were λ = 561 nm for TRITC and λ = 491 nm for GFP and fluorescein. The excitation objective was a 10×, 0.3 NA Plan Fluor from Nikon, while the detection objective was a 10×, 0.6 NA Plan Apochromat from Olympus. The latter had a correction collar for the refractive index of the immersion solution, ranging from 1.33 to 1.52. The samples were placed in a quartz cuvette containing the mounting medium (63% TDE/PBS) and placed in a custom-made chamber filled with the mounting medium. The samples were mounted on a motorized *x*-, *y*-, *z*-, *Θ*-stage (M-122.2DD and M-116.DG, Physik Instrumente, Germany), which allowed free 3-D motion and rotation. Stacks were acquired with a *z*-step of 2 μm and and an *xy* resolution resulting from the setup configuration of 0.65 μm, with a field of view of 1.3×1.3 mm. The microscope was controlled via custom written LabVIEW code (National Instruments), which coordinated the galvo scanners, the rolling shutter, and the stack acquisition.

### Evaluation of morphological changes

To expose the brain for *in vivo* two-photon acquisition, a craniotomy^36, 37^ was performed on a single adult C57BL/6 mouse. Mice were deeply anesthetized by intraperitoneal injection of zoletil (50 mg/kg) and xylazine (9 mg/kg). A small dose of dexamethasone (0.04 ml at 2 mg/ml) was administered to minimize swelling at the surgical site. Using a dental drill, a circular portion of the skull (5 mm in diameter) above the motor cortex was removed. The exposed region was covered with a cover glass and sealed with dental cement. The experimental protocol was designed in accordance with the rules of the Italian Ministry of Health.

*In vivo* brain-vasculature staining was performed by injecting 0.2 ml of Texas red Dextran (140 μg/ml, in saline solution) into the tail vein. Two-photon acquisition was performed through the brain window soon after *in vivo* vasculature labelling.

*Ex vivo* two-photon imaging of the same brain region was performed on the dissected brain after staining with the protocol described in Tsai et al.^21^ Large vessels were used as a reference to identify the same area.

After clearing, whole-brain images were acquired with LSFM and the selected region was retrieved from the whole-brain dataset.

Vessel diameter was determined with ImageJ/Fiji software (http://fiji.sc/Fiji). Blood vessel length was measured by manually tracing selected vessels with the Filament Editor function of Amira 5.3 software (FEI Visualization Sciences Group, Oregon, USA).

### Image segmentation

We manually and automatically segmented the stacks acquired via TPFM and LSFM. For both TPFM and LSFM derived images, we used Amira 5.3 software (FEI Visualization Sciences Group, Oregon, USA) for the manual segmentation and an algorithm based on the Markov random fields^24^ for the automatic segmentation. We applied the co-localization tool in ImageJ/Fiji (JACoP plug-in) to compute true positive, false positive, and false negative rates of the automatic segmentation with respect to the manual segmentation. From these values, we were able to estimate the precision and recall of the automatic segmentation.

### Images stitching and 3D renderings

LSFM produced a series of 3D stacks, each one covering an area of 1.3×1.3 mm, with regions of superimpositions. To achieve a 3D image of whole specimens from raw data, the TeraStitcher tool^38^, a sofware capable of dealing with teravovel sized images, was used. 3D renderings of stitched images of whole mouse brains were produced from downsampled files using the Amira Voltex function of Amira 5.3 software (FEI Visualization Sciences Group, Oregon, USA).

### Data analysis

Graphs and data analysis were done with OriginPro 9.0 (OriginLab Corporation). All data values are given as means ± SD. Data were analysed using one-way ANOVA with Bonferroni’s test for multiple comparisons. *P* values of ≤ 0.05 were considered significant. Image stacks were analysed using both Fiji (http://fiji.sc/Fiji) and Amira 5.3 software (FEI Visualization Sciences Group, Oregon, USA).

## Acknowledgements

This project received funding from the European Union’s H2020 research and innovation programme under grant agreements No. 720270 (Human Brain Project) and 654148 (Laserlab-Europe), and from the EU programme H2020 EXCELLENT SCIENCE - European Research Council (ERC) under grant agreement ID n. 692943 (BrainBIT). The project have also been supported by the Italian Ministry for Education, University, and Research in the framework of the Flagship Project NanoMAX and of Eurobioimaging Italian Nodes (ESFRI research infrastructure), and by “Ente Cassa di Risparmio di Firenze” (private foundation).

## Author contribution

A.P.D.G., L.Sil and L. Sac. Planned the experiments; A.P.D.G. performed the vascular stainings, prepared the clarified samples and execute the manual segmentations. A.T. and P.F. performed the automatic segmentations. I.C. established the clarification procedure. M.C.M., L.Sil. and A.P.D.G. performed LSFM imaging. A.P.D.G. imaged the samples using TPFM. A.L.A.M. performed the craniotomy and implantation of the cranial window. F.S.P. supervised the project. A.P.D.G. made the figures and wrote the paper with the input of all the other authors.

## FIGURE LEGENDS

**Supplementary video 1**

Whole-brain 3D rendering of the vasculature network. Image obtained with Amira 5.3 software from downsampled data. Stacks were stitched using TeraStitcher software^38^.

**Supplementary video 2**

3D rendering of the vasculature from 1.3×1.3×2 mm ventral brain region at the correspondence of the right middle cerebral artery sprouting from the Circle of Willis. Image obtained with Amira 5.3 software from downsampled LSFM image stack.

**Supplementary video 3**

3D rendering of 1.3×1.3×2 mm from downsampled image stack showing GFP-expressing neurons under the *thy1* promoter. The *Thy1*-GFP-M brain sample was treated for vasculature staining and clearing as described. Image obtained with Amira 5.3 software from downsampled LSFM image stack.

**Supplementary video 4**

3D rendering showing blood vessels from the same portion of brain showed in supplementary video 3. Image obtained with Amira 5.3 software from downsampled LSFM image stack.

